# Identification of LMAN1 and SURF4 dependent secretory cargoes

**DOI:** 10.1101/2023.04.06.535922

**Authors:** Vi T. Tang, Prabhodh S. Abbineni, Felipe da Veiga Leprevost, Venkatesha Basrur, Brian T. Emmer, Alexey I. Nesvizhskii, David Ginsburg

## Abstract

Most proteins secreted into the extracellular space are first recruited from the endoplasmic reticulum into coat protein complex II (COPII)-coated vesicles or tubules that facilitate their transport to the Golgi apparatus. Although several secreted proteins have been shown to be actively recruited into COPII vesicles/tubules by the cargo receptors LMAN1 and SURF4, the full cargo repertoire of these receptors is unknown. We now report mass spectrometry analysis of conditioned media and cell lysates from HuH7 cells CRISPR targeted to inactivate the *LMAN1* or *SURF4* gene. We found that LMAN1 has limited clients in HuH7 cells whereas SURF4 traffics a broad range of cargoes. Analysis of putative SURF4 cargoes suggests that cargo recognition is governed by complex mechanisms rather than interaction with a universal binding motif.

## Introduction

Approximately a third of the proteins encoded by the mammalian genome are co-translationally inserted into the endoplasmic reticulum (ER), from where they are subsequently trafficked to the plasma membrane, various intracellular organelles, or secreted into the extracellular space (1-3). Properly folded proteins destined for secretion are transported from the ER to the Golgi via coat protein complex II (COPII) coated vesicles or tubular structures (4-6). Entry into COPII vesicles/tubules is thought to occur passively via bulk flow or through active recruitment and concentration (7). Transmembrane cargoes can interact directly with the cargo selective COPII component SEC24 on the cytoplasmic face of the ER, whereas soluble proteins restricted to the ER lumen require interaction with a membrane-spanning intermediary to bridge this interaction. However, relatively few such cargo receptors have been discovered in mammalian cells to date and the cargo receptor-dependence for most secreted proteins remains unclear.

LMAN1, also known as ERGIC-53, is a 53 kDa protein that localizes to the ER and ER-Golgi intermediate compartment (ERGIC) (8). LMAN1 has been shown to function as a cargo receptor for coagulation factors V (F5) and VIII (F8) (9,10), alpha-1-antitrypsin (SERPINA1) (11), Mac-2 binding protein (Mac-2BP) (12), matrix metalloproteinase-9 (MMP9) (13), cathepsin C (CTSC) (14), cathepsin Z (CTSZ) (15), and membrane protein γ-aminobutyric acid type A receptors (GABAARs) (16). *In vivo* studies in mice have confirmed the dependence of F5, F8, and SERPINA1, but not CTSC and CTSZ, on LMAN1 for secretion (17). No common LMAN1 binding motif has been identified. It is unclear whether there are other cargoes beyond those listed above that require LMAN1 for efficient secretion from the ER.

Another recently identified mammalian cargo receptor, SURF4, is a 29 kDa protein with multiple transmembrane domains that also localizes to the ER and ERGIC (18). SURF4 is highly conserved, with homologs in yeast (Erv29p), *C. elegans* (SFT-4), and *Drosophila* (19). Erv29p facilitates the secretion of yeast pro-α-factor, carboxypeptidase Y, and proteinase A (20-22). Multiple SURF4 cargoes in mammals have been identified to date, including PCSK9 (23-25), apolipoprotein B (APOB) (25-28), growth hormone (29), dentin sialophosphoprotein (DSPP) (29), amelogenin (29), erythropoietin (30), pathogenic SERPINA1 polymers (31), sonic hedgehog (32), proinsulin (33), and the lysosomal proteins progranulin and prosaposin (34). Two SURF4 binding motifs on cargoes have been proposed, including an ER-ESCAPE tripeptide motif immediately downstream of the signal peptide sequence (29) and a Cardin-Weintraub motif (32,35), though not all of the putative cargoes listed above carry one of these motifs, suggesting the presence of additional determinants for recognition by SURF4 (24).

Previous attempts to define a comprehensive cargo repertoire for LMAN1 and SURF4 have revealed additional putative cargoes. Using an *in vitro* vesicle formation assay and label-free mass spectrometry in cells depleted for LMAN1 or SURF4, Huang et al (36) described 4 and 17 novel cargoes, respectively for these receptors. Using SILAC labeling and mass spectrometry analysis of conditioned media following SURF4 knockdown, Gomez-Navarro and colleagues (24) likewise identified 10 proteins in HEK293 cells and 18 proteins in HuH7 cells as potential SURF4 cargoes.

We and others have reported a mass spectrometry approach to identify secreted proteins by quantifying protein abundance in both conditioned media and cell lysates from cultured cells (37-39). We found that calculation of media to lysate (M/L) ratios (rather than using unadjusted protein abundance in the media alone) greatly improved the sensitivity and specificity for detecting secreted proteins (39). We now report the application of this approach to determine the cargo repertoire of LMAN1 and SURF4 in HuH7 cells.

## Materials and Methods

### Cell culture

HuH7 cells (40) were cultured in DMEM supplemented with Glutamax (ThermoFisher Scientific, Waltham MA, 10569-044), 10% fetal bovine serum (MilliporeSigma, Burlington MA, F8067), and penicillin/streptomycin (ThermoFisher Scientific, Waltham MA, 15140-122). Cells were passaged every 3-4 days and maintained between 20-80% confluence.

### CRISPR mediated inactivation of LMAN1 and SURF4

On day 0, cells were seeded at 20% confluence and infected with lentivirus generated from pLentiCRISPRv2 (Addgene #52961, a gift from Feng Zhang (41)) engineered to deliver Cas9 and a gRNA targeting *LMAN1* (CCCCTTACACTATAGTGACG), *SURF4* (TCCGAGCTGCATGTACTGTT), or a nontargeting gRNA (GTTCATTTCCAAGTCCGCTG) as previously described (42). On day 1, the medium was exchanged and 1 µg/ml puromycin (MilliporeSigma, Burlington MA, P8833) was added for 48 hours. Following selection, surviving cells were passaged every 3-4 days until day 14 to allow for gene editing and protein turnover. On day 14, cells were washed three times with phosphate-buffered saline (PBS, ThermoFisher Scientific, Waltham MA, 10010-023) and switched to serum-free, phenol red-free DMEM prewarmed to 37ºC (ThermoFisher Scientific, Waltham MA, 31053-036) for 12 hours.

### Conditioned media and cell lysate collection

Conditioned media and cell lysates were harvested as previously described (39). Briefly, conditioned media were collected from the cell culture dish and centrifuged at 2,500g at 4ºC for 15 minutes to remove cell debris. The supernatant was then ultracentrifuged at 120,000g at 4ºC for 90 minutes to remove exosomes (43), and concentrated using a 3 kDa molecular weight cutoff concentrator (MilliporeSigma, Burlington MA, UFC900324). Cell lysates were collected in 2 ml of RIPA buffer (Thermo Scientific, 89900) containing a protease inhibitor cocktail (cOmplete™, Mini Protease Inhibitor Cocktail, Roche, Basel, Switzerland, 11836153001). Cell suspensions were sonicated, rotated end-over-end for 1 hour, and centrifuged at 21,000g at 4ºC for 45 minutes. Supernatants were then transferred to new Eppendorf tubes. Protein concentration in the conditioned media and lysates were determined by DC protein assay (Bio-Rad, Hercules CA, 500-011).

### Immunoblotting

Cell lysates (10 μg per sample) collected as above were resolved in a 4-20% Tris-glycine gel as previously described (25). Proteins were detected with antibodies against LMAN1 (Abcam, Cambridge UK, ab125006, 1:1000) or GAPDH (Abcam, Cambridge UK, ab181602, 1:10000).

### Mass spectrometry, protein identification, and protein quantification

Mass spectrometry, protein identification, and protein quantification were performed as previously described (39). Briefly, 75 μg of each lysate and 75 μg of each medium sample were proteolyzed, labeled with tandem mass tags (TMT) 10-plex according to manufacturer’s protocol, and subjected to liquid chromatography-mass spectrometry analysis. Raw mass spectrometry files were converted into open mzML format and were analyzed using the FragPipe (https://fragpipe.nesvilab.org/) computational platform (44-46) with the default TMT10-MS3 workflow. For proteins that were identified and quantified in both the media and lysate fractions, a M/L ratio was calculated using absolute intensity values. Signal peptide and transmembrane domain annotations were obtained from the UniProt database (47).

### Statistical analyses

The limma statistical package was used for comparison of protein abundance or M/L ratios between cells treated with a nontargeting, *LMAN1*-targeting, SURF4-targeting gRNA, using a log2-transformed protein abundance or M/L ratios as input (48). P-values were adjusted for multiple hypothesis testing using the Benjamini & Hochberg method. An adjusted p-value (q-value) of 0.05 or less was considered statistically significant.

### Analysis of ER-ESCAPE motifs

The human proteome reference database was downloaded from Uniprot (47). Proteins were filtered for the presence of a signal peptide and absence of a transmembrane domain(s). The tripeptide motif as proposed by Yin et al (29) was extracted from the protein sequence. For proteins with a conventional signal peptide, the tripeptide motif is defined as the first three amino acid residues downstream of the annotated signal peptide sequence. For proteins with a propeptide domain, the tripeptide motif is defined as the first three amino acid residues downstream of the propeptide cleavage site. For rare cases of proteins with an uncleaved signal peptide (based on Uniprot annotation), the tripeptide motif is assigned as the first three amino acid residues of the protein. Each residue within the tripeptide motif was classified based on the classification system proposed by Yin et al (29). For this analysis, SURF4 cargoes include putative SURF4 cargoes identified in this study as well as all previously reported cargoes.

## Results

### Identification of bona fide secreted proteins by analysis of cell lysates and conditioned media

To identify secreted proteins that depend on either LMAN1 or SURF4 for secretion, *LMAN1* and *SURF4* deficient HuH7 cells were generated by CRISPR mediated gene editing. We then collected conditioned media and cell lysates from cells receiving *LMAN1*-targeting, *SURF4*-targeting, or nontargeting (NT) gRNA for analysis by TMT mass spectrometry (n=3 per group) (**Figure 1A**).

**Figure 1:**
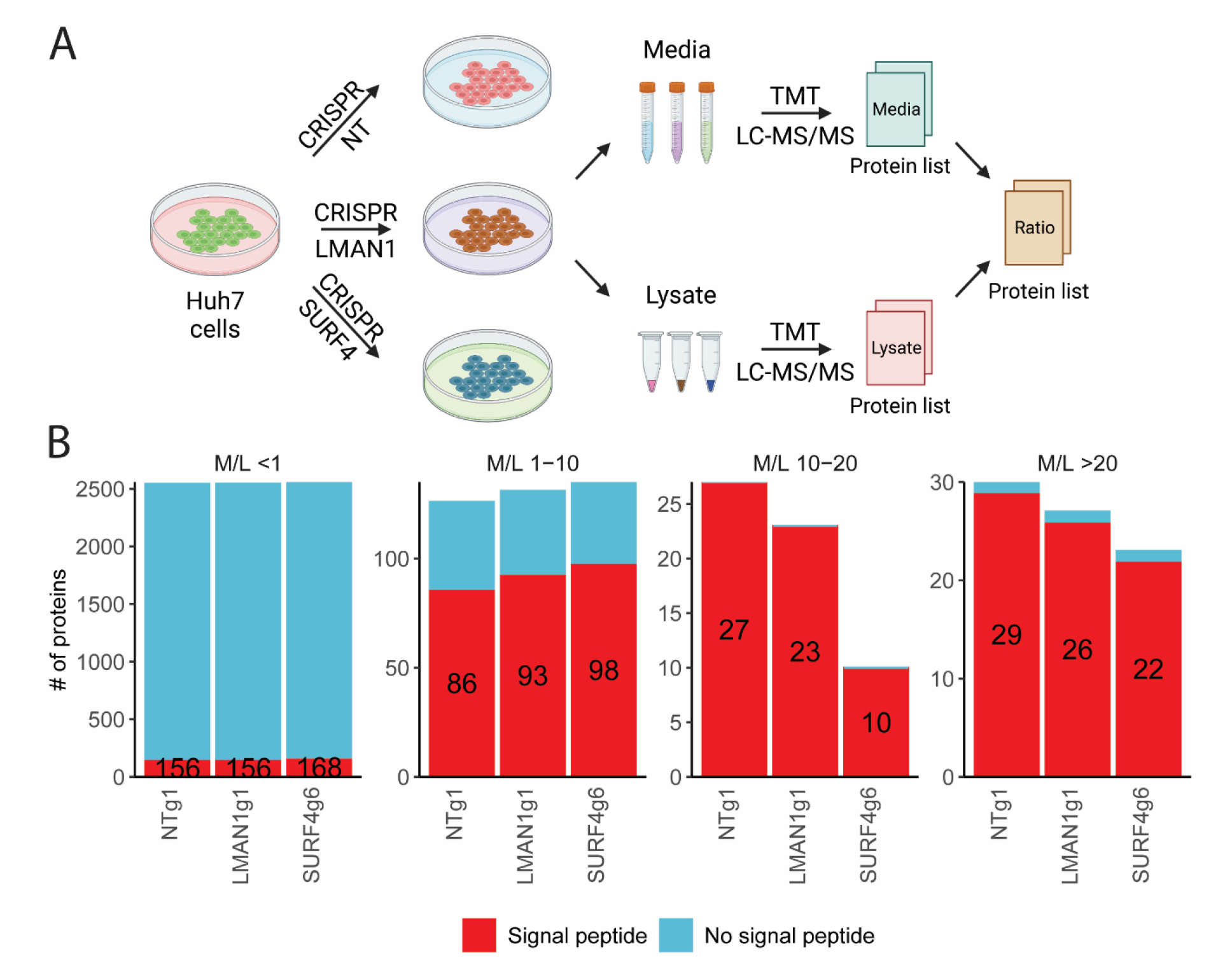
Identifying proteins dependent on LMAN1 or SURF4 for efficient secretion. **(A)** Experimental design to identify LMAN1 and SURF4 cargoes in the human hepatoma cell line (HuH7). HuH7 cells were infected with lentiviruses delivering CRISPR/Cas9 and guide RNAs targeting either *LMAN1, SURF4*, or a nontargeting (NT) control. Following selection, cells were cultured for 2 weeks before being switched to serum free media for 12 hours. Conditioned media and cell lysates were collected for protein identification and quantification by liquid chromatography (LC) followed by tandem mass tag (TMT) mass spectrometry (MS). A protein abundance ratio was calculated for each protein that was detected in both the media and lysate fractions as described in Methods. **(B)** Number of proteins with (red) and without (blue) a signal peptide identified in different M/L fractions. The number of signal peptide-containing proteins in each bar is indicated.

Across all three sample groups, we identified and quantified 5858 and 2947 proteins in the lysate and media fractions, respectively, of which 2726 proteins were identified in both fractions (**Figure S1A** and **Table S1-2**). Consistent with our previous observations (39), the majority of identified proteins lack a signal peptide, with increasing M/L ratio correlating with an increased proportion of proteins carrying a signal peptide, consistent with the expected enrichment for secretory proteins in the media relative to the cell lysates (**Figure S1B**). Comparisons between NT, LMAN1, and SURF4 samples revealed that there were fewer proteins with an M/L ratio greater than 10 in LMAN1- and SURF4-deficient samples (**Figure 1B**).

### Few proteins in HuH7 cells depend on LMAN1 for secretion

To identify secretory proteins that require LMAN1 for their secretion, we compared the M/L ratios of proteins collected from cells treated with LMAN1-targeting or NT gRNA. As shown in **Figure 2A**, the only protein demonstrating a significantly different M/L ratio following *LMAN1* deletion was MCFD2, a 16 kDa soluble ER luminal protein that forms a complex with LMAN1 and acts as cofactor for the secretion of factors V and VIII. MCFD2 lacks an ER retrieval motif and its retention in the ER and ERGIC depends entirely on its interaction with LMAN1 (49). The increased M/L ratio for MCFD2 in LMAN1-deficient cells (**Figure 2A** and **Table S3**), resulting from a decreased intracellular (**Figure S2A** and **Table S4**) and increased extracellular abundance (**Figure S2B** and **Table S5**), is therefore consistent with its release from the ER in the absence of LMAN1 (36,49).

**Figure 2.**
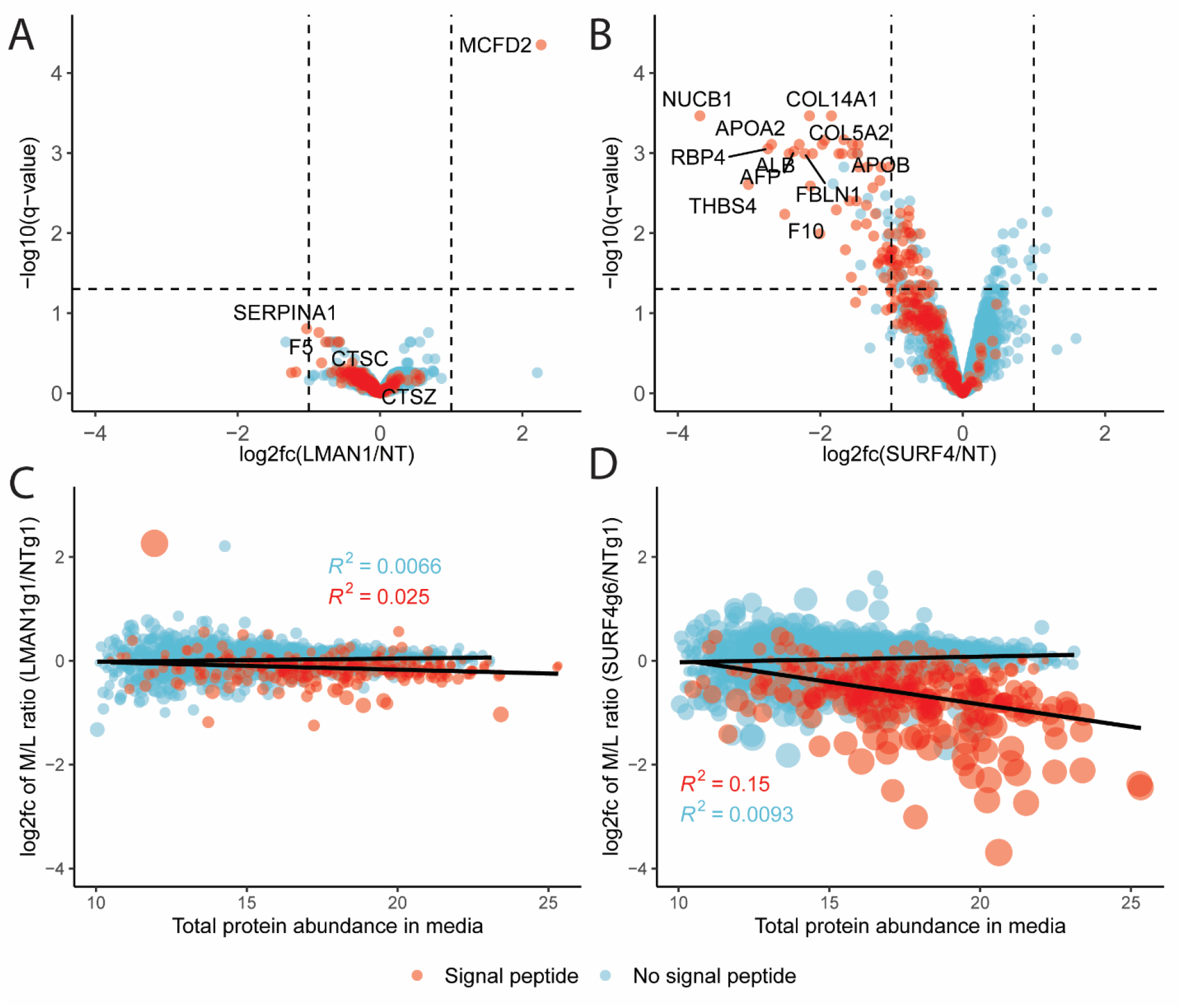
Differential effects on protein secretion in HuH7 cells following LMAN1 or SURF4 deletion. **(A-B)** Volcano plots comparing protein M/L ratios in LMAN1 **(A)** or SURF4 **(B)** deficient cells with those in controls. The log2 fold change (log2fc) and statistical significance are plotted on the x and y-axis, respectively. Proteins with a signal peptide are colored in red and proteins without a signal peptide are colored in blue. Dashed vertical lines represent the log2fc of 1 and −1. Dashed horizontal lines represent the log10(q-value of 0.05). **(C-D)** Comparison of total abundance (estimation from Fragpipe, see Methods) in the media with LMAN1 **(C)** or SURF4 **(D)** dependency. Trend lines represent linear regression. Dot sizes are proportional to log10(q-value). R^2^ values in blue and red are for proteins without and with an annotated signal peptide, respectively.

Consistent with previously reported *in vivo* data in mice, We also found that genetic deletion of LMAN1 in HuH7 cells resulted in intracellular accumulation of SERPINA1 (50) (**Figure S2A**) without a statistically significant reduction in media abundance (**Figure S2B**) or M/L ratio (**Figure 2A**). Though a trend was observed consistent with decreased secretion of F5, a known LMAN1 cargo, these changes did not reach statistical significance. Taken together, these data suggest that LMAN1 traffics a small number of secretory cargoes in HuH7 cells. Though we cannot exclude technical limitations including incomplete LMAN1 deletion, the marked LMAN1 depletion observed in LMAN1-deficient cell lysates by mass spectrometry (**Figure S2A**) and immunoblotting (**Figure S3**) suggests that this latter explanation is unlikely.

### A wide range of proteins depend on SURF4 for secretion in HuH7 cells

In contrast to LMAN1, SURF4 disruption in HuH7 cells caused a significant decrease in the M/L ratio for numerous signal peptide-containing proteins (log2(fc) < −1 and adjust p-value < 0.05). For most proteins, this change was associated with both an increase in lysate abundance and a decrease in media abundance (**Figure S4** and **Table S6-7**). Affected proteins include known SURF4 cargoes such as APOB, APOA1, and APOA2 (27,28), several putative cargoes identified in other mass spectrometry based studies (24,36), as well as a number of potentially novel SURF4 cargoes (**Figure 2B** and **Table S8**). Though PCSK9 has been demonstrated to be dependent on SURF4 for efficient secretion *in vivo* (25) and in HEK293T cells *in vitro* (23). We did not identify PCSK9 as a SURF4 cargo in HuH7 cells, consistent with a previous report (51). Seven out of 2726 proteins in our dataset were among the other known SURF4 cargoes from previously published studies, four of which showed significantly decreased M/L ratio. Comparison of protein abundance in the media with the effect of cargo receptor deletion suggested a greater dependency of highly abundant proteins on SURF4, but not LMAN1, for their secretion (**Figure 2C-D**). A comparison of the putative SURF4 cargoes identified here with those in two other recent reports (24,36) identified four secreted proteins shared between all datasets (**Figure 3A** and **Table S9**) and ten between two or more datasets (**Figure 3A** and **Figure 3C**).

**Figure 3.**
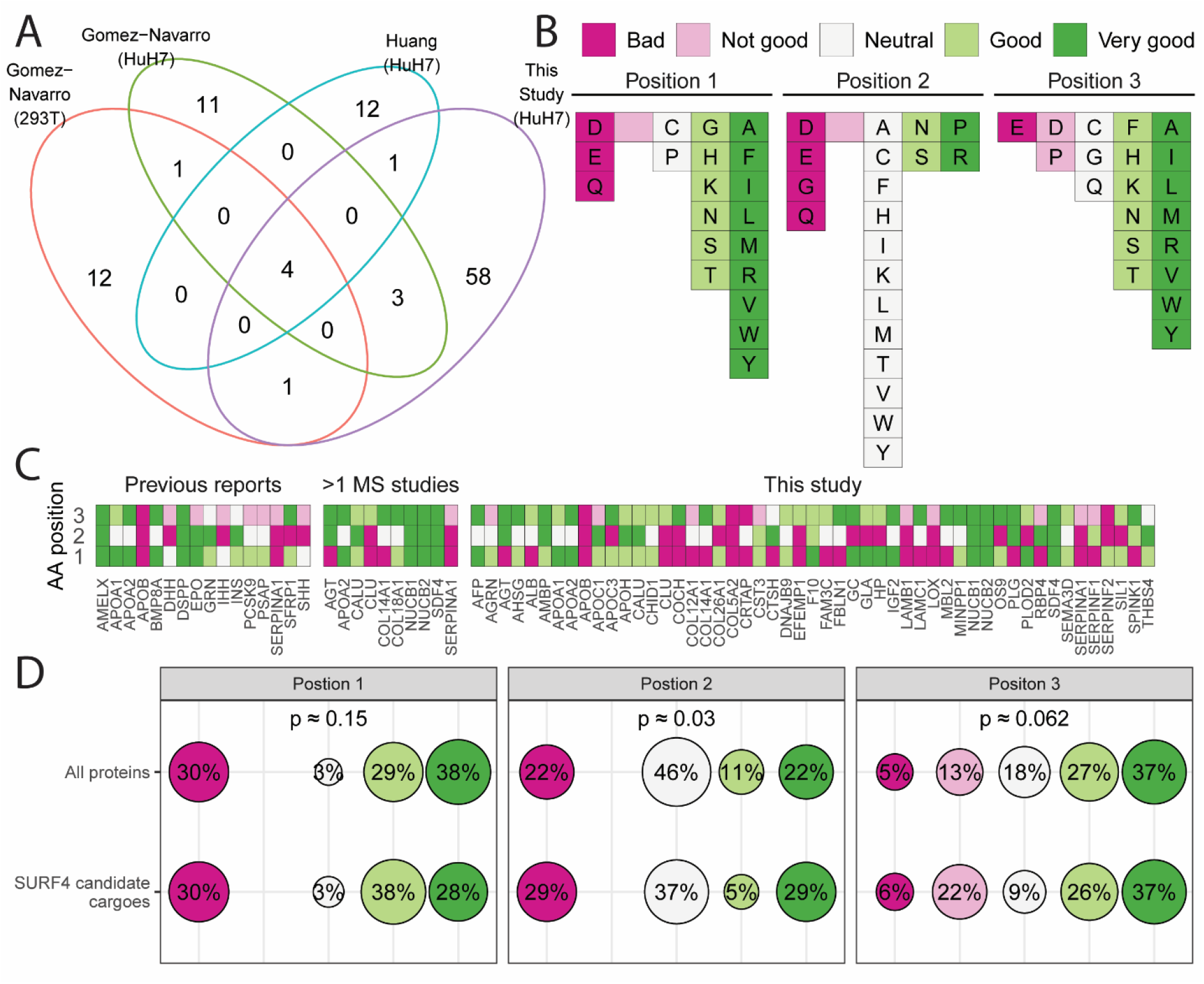
Most SURF4 cargoes do not contain an ER-ESCAPE motif (29). **(A)** Venn diagram of putative SURF4 cargoes that were identified in the current study, or previously reported by Gomez-Navarro et al (24) in HEK293T and HuH7 cells following analyses of conditioned media, or by Huang et al (36) using an *in vitro* COPII vesicle formation assay. **(B)** A classification system for the three amino acid residues immediate downstream of the signal peptide cleavage site (ER-ESCAPE tripeptide motif) proposed by Yin et al (29). **(C)** Tripeptide motifs in previously reported SURF4 cargoes (23-34), in cargoes that were identified in more than one MS-based datasets, and in this study color-coded according to (B). **(D)** Percentage of each amino acid class (as described in (B)) at each position in the tripeptide motif for all secreted proteins with an annotated signal peptide in the proteome (n = 2110, see Methods) and for SURF4 candidate cargoes (n = 86). P-values were obtained from Fisher’s exact test.

### Most SURF4-dependent secreted proteins lack an ER-ESCAPE motif

Previously, Yin et al (29) proposed a tripeptide motif (ER-ESCAPE) as being responsible for the recognition and recruitment of SURF4 cargoes. This motif is located immediately downstream of the signal peptide and is characterized by a proline residue flanked on either side by a hydrophobic amino acid (**Figure 3B**). Qualitative analysis of candidate SURF4 cargoes identified in our study and in previous studies revealed a wide range of sequences at the predicted site of the ER-ESCAPE motif, including some that fit the motif well (NUCB1, NUCB2, and SDF4) and others with no apparent overlap (COL5A2 and APOB) (25-28) (**Figure 3C** and **Table S10**).

To determine whether the ER-ESCAPE motif was significantly enriched among SURF4 cargoes, we compared the tripeptide sequences from the full set of candidate SURF4 cargoes in relation to all proteins in the human proteome with an annotated signal peptide (**Table S11-12**). As shown in **Figure 3D**, there are no significant differences in the distribution of each amino acid class within the tripeptide motif between SURF4 candidate cargoes that have been reported to date and all secreted proteins with an annotated signal peptide in the proteome. This analysis suggests that the ER-ESCAPE motif may only be relevant for a limited subset of SURF4 cargoes.

Recent studies have also suggested a key role for the Cardin-Weintraub (CW) motif [K/R][K/R][K/R]XX[K/R][K/R] in recognition by SURF4 for cargoes such as the hedgehog family (32,35). Analysis of the human proteome identified 267 proteins carrying a CW motif among 7327 proteins that carry a signal peptide or a transmembrane domain. Among the 115 SURF4 candidate cargoes identified by our and previous studies (**Figure 3C**), seven carry a CW motif though only one of these seven proteins was identified in the current study (**Table S13**).

## Discussion

In this study, we applied a broad, whole proteome approach (39) to profile the cargo repertoires for two well-characterized cargo receptors, LMAN1 and SURF4, in the human hepatocellular carcinoma cell line, HuH7. Though only a limited set of LMAN1-dependent cargoes were identified in HuH7 cells, SURF4 facilitated secretion for a broad range of proteins. A subset of putative SURF4 cargoes carry the previously described ER-ESCAPE (29) or CW motifs (32,35) potentially mediating interaction with SURF4. However, most do not, suggesting a more complex mechanism governing SURF4 cargo recognition.

While several established LMAN1 cargoes, including F5 and SERPINA1, were detected in our dataset, with a trend toward reduced M/L ratio in LMAN1 deleted cells, these changes did not reach statistical significance. These findings suggest limited statistical power, potentially due to only partial secretion blockade in LMAN1-deficient HuH7 cells, and/or overlap with other cargo receptors. Indeed, humans and mice with completely deficient for LMAN1 exhibit incomplete reduction in plasma levels for 2 key LMAN1 cargoes, F5 and F8, to only ∼10% (humans) and 50% (mice) of those in wild-type controls (50,52). Similarly, levels for the LMAN1 cargo SERPINA1 are unchanged in the plasma of *Lman1*^*-/-*^ mice, though modest accumulation is observed in hepatocytes (50). Although the lysosomal proteins CTSC and CTSZ have also been proposed as cargoes for LMAN1 (14,15), our experiment approach would not be expected to distinguish protein accumulation in different cellular compartments (i.e. the ER vs. the lysosome). Taken together, these data suggest that LMAN1 only traffics a small subset of proteins in HuH7 cells.

In contrast to LMAN1, depletion of SURF4 significantly impeded secretion of 52 proteins, suggesting a much broader cargo repertoire for SURF4 than for LMAN1. SERPINA1, a previously reported LMAN1 cargo (11), also exhibited reduced M/L ratio following SURF4 deletion, suggesting dependence on SURF4 as well as LMAN1 for efficient secretion. These findings are consistent with a recent report demonstrating a role for SURF4 in the secretion of pathogenic SERPINA1 polymers as well as SERPINA1 monomers, albeit to a lesser extent than LMAN1 (24,31). The large difference in the number of potential clients for LMAN1 and SURF4 could also help explain the normal development and only mild bleeding defect observed in LMAN1 deficient mice and human (50,52) in contrast to the early embryonic lethality of *Surf4*^*-/-*^ mice (53) and the lack of human disorders associated with mutations in *SURF4*.

Though the role of the ER-ESCAPE motif proposed by Yin et al (29) in the efficient SURF4-mediated trafficking of PCSK9 and NUCB1 has been confirmed by Gomez-Navarro and colleagues (24), our analysis suggests a more complex process for SURF4 cargo selection. We failed to confirm a general enrichment for the ER-ESCAPE motif among the broader repertoire of potential SURF4 cargoes. However, we cannot exclude the possibility that the tripeptide motif is shifted from the expected starting position in the majority of cargoes (24). It is also possible that SURF4 indirectly interacts with cargoes via a cofactor with each cofactor having a different recognition motif. Lastly, the broad range of proteins that rely on SURF4 for secretion also suggests the possibility that SURF4 could function as a general mediator of ER-Golgi transport, promoting secretion in a way other than directly binding to its cargoes. For example, a recent study reported that SURF4 regulates entrance of secretory proteins into a tubular network lacking LMAN1 and extending from the ER, which expedites proteins delivery to the Golgi, suggesting a distinct SURF4 trafficking route (54).

In this report, we demonstrated the analysis of protein abundance in both cell lysates and condition media to identify secreted proteins that depend on LMAN1 and SURF4 for secretion. This approach could also be applied to other cargo receptors and cell types. In addition, a similar strategy could also be extended to study changes in cell secretomes resulting from disruptions in the conventional and unconventional secretory pathway as well as in response to external stimuli.

## Acknowledgements

This work was supported by grants from the National Institutes of Health (R35HL135793 to DG, R01GM094231 to AIN, and K99GM141268 to PSA). VTT was supported by fellowships from the American Heart Association Predoctoral Fellowship (20PRE35210706) and the University of Michigan Rackham Graduate School. DG is a Howard Hughes Medical Institute Investigator.

---

**Figure S1.**
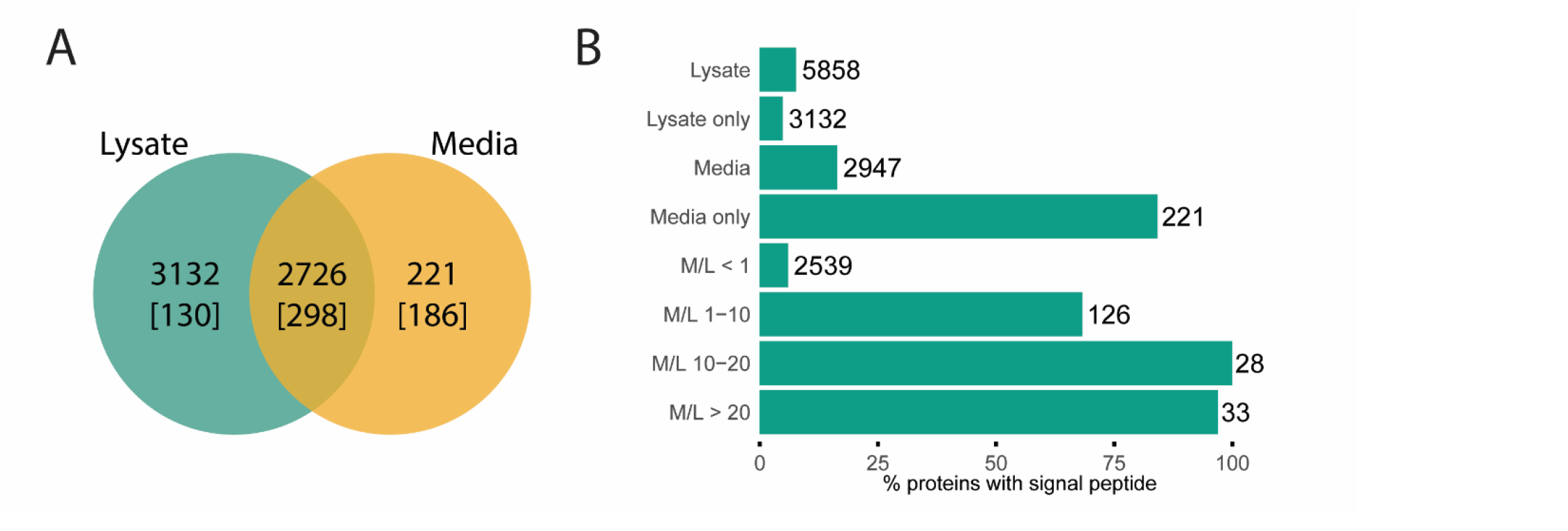
Comparison of protein abundance in the cell media and lysates improves identification of secreted proteins. **(A)** Venn diagram of proteins detected in conditioned media, cell lysates, or both fractions across all samples (LMAN1-defient, SURF4-deficient, and NT-treated cells, n = 3 per group). Numbers in the bracket represent the number of proteins carrying an annotated signal peptide in each group. **(B)** Proteins were separated into discrete media/lysate (M/L) ratio intervals and the percentage of proteins carrying a signal peptide was calculated. Numbers at the end of each bar indicate the number of proteins in each bin.

**Figure S2.**
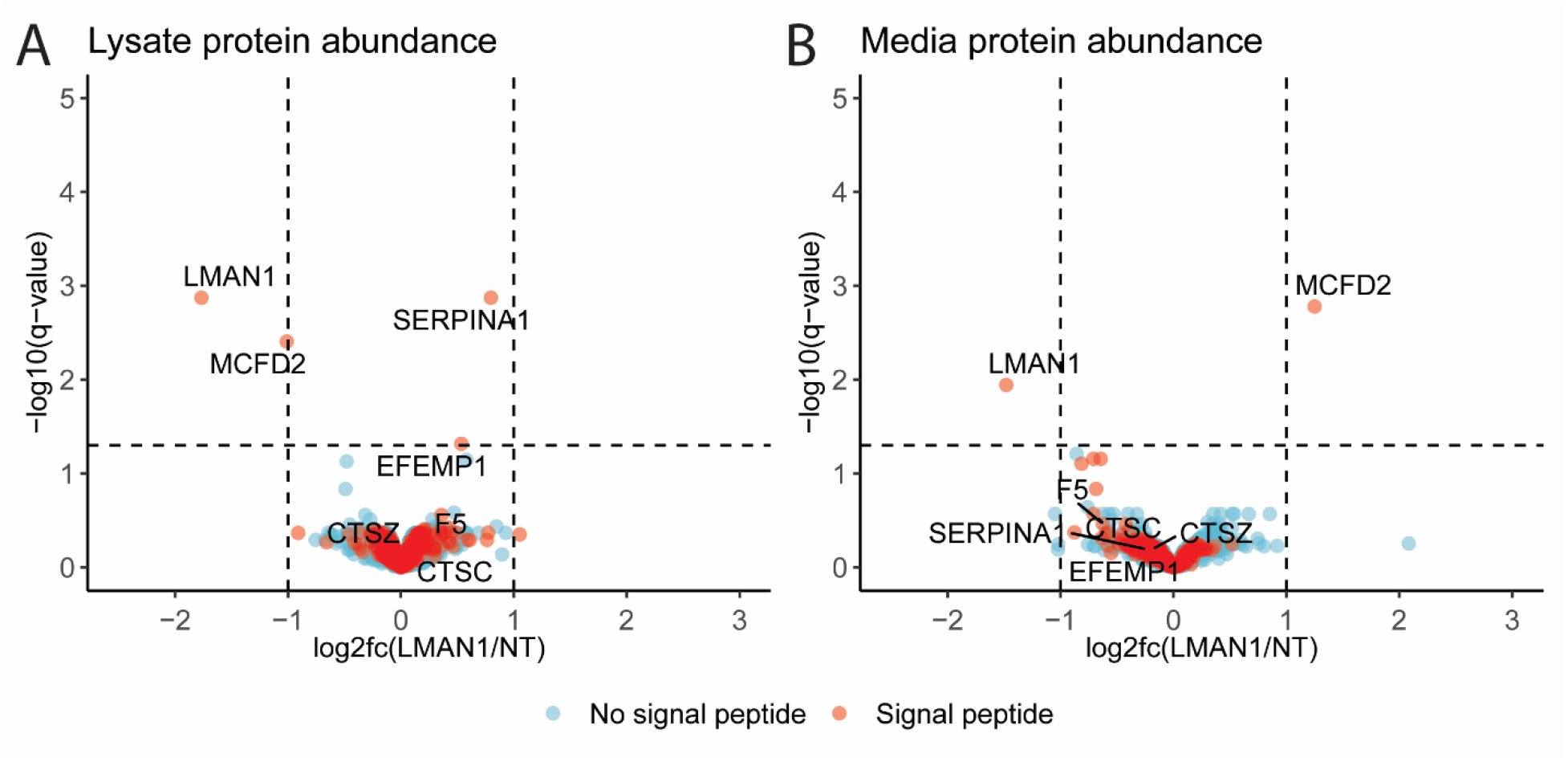
Comparison of proteins abundance in the lysates and media between LMAN1 deficient and control cells. Volcano plots representing changes in protein abundance in the cell lysates **(A)** or conditioned media **(B)**. The log2 fold change (log2fc) and statistical significance are plotted on the x and y-axis, respectively. Proteins with a signal peptide are colored in red and proteins without a signal peptide are colored in blue. Dashed vertical lines represent the log2fc of 1 and −1. Dashed horizontal lines represent the log10 (q-value of 0.05).

**Figure S3.**
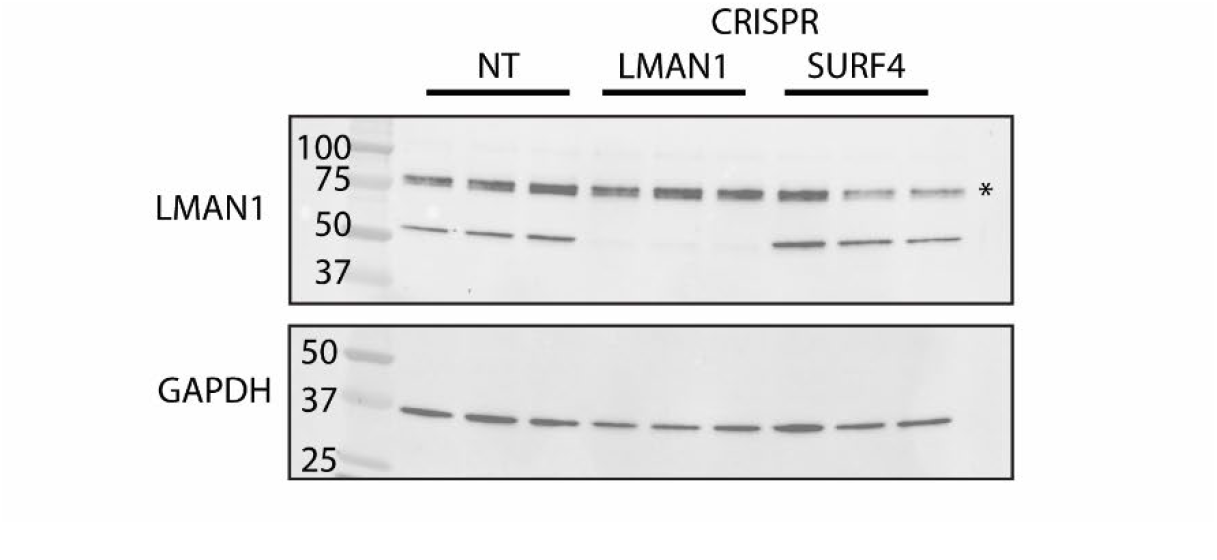
Efficient depletion of LMAN1 in cells treated with *LMAN1* targeting gRNA. Immunoblots of cell lysates collected from controls (NT) and *LMAN1* or *SURF4* deleted cells. Numbers represent protein molecular weights in kDa. Asterisk indicates nonspecific binding of LMAN1 antibody.

**Figure S4.**
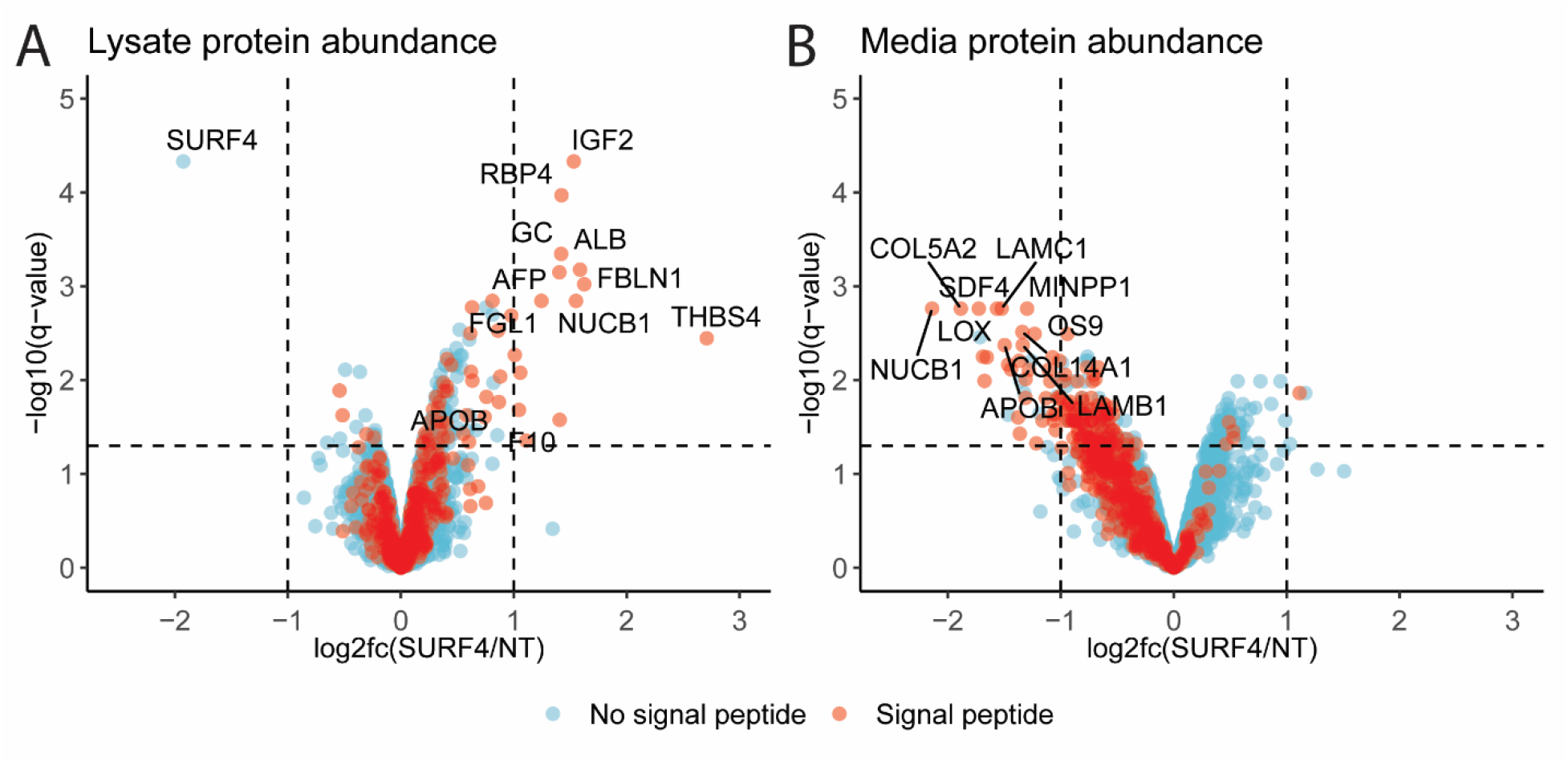
Comparison of proteins abundance in the lysates and media between SURF4 deficient and control cells. Volcano plots representing changes in protein abundance in the cell lysates **(A)** or conditioned media **(B)**. The log2 fold change (log2fc) and statistical significance are plotted on the x and y-axis, respectively. Proteins with a signal peptide are colored in red and proteins without a signal peptide are colored in blue. Dashed vertical lines represent the log2fc of 1 and −1. Dashed horizontal lines represent the log10 (q-value of 0.05).

## References

1. Uhlen, M., Fagerberg, L., Hallstrom, B. M., Lindskog, C., Oksvold, P., Mardinoglu, A., Sivertsson, A., Kampf, C., Sjostedt, E., Asplund, A., Olsson, I., Edlund, K., Lundberg, E., Navani, S., Szigyarto, C. A., Odeberg, J., Djureinovic, D., Takanen, J. O., Hober, S., Alm, T., Edqvist, P. H., Berling, H., Tegel, H., Mulder, J., Rockberg, J., Nilsson, P., Schwenk, J. M., Hamsten, M., von Feilitzen, K., Forsberg, M., Persson, L., Johansson, F., Zwahlen, M., von Heijne, G., Nielsen, J., and Ponten, F. (2015) Tissue-based map of the human proteome. Science 347, 1260419

2. Uhlen, M., Karlsson, M. J., Hober, A., Svensson, A. S., Scheffel, J., Kotol, D., Zhong, W., Tebani, A., Strandberg, L., Edfors, F., Sjostedt, E., Mulder, J., Mardinoglu, A., Berling, A., Ekblad, S., Dannemeyer, M., Kanje, S., Rockberg, J., Lundqvist, M., Malm, M., Volk, A. L., Nilsson, P., Manberg, A., Dodig-Crnkovic, T., Pin, E., Zwahlen, M., Oksvold, P., von Feilitzen, K., Haussler, R. S., Hong, M. G., Lindskog, C., Ponten, F., Katona, B., Vuu, J., Lindstrom, E., Nielsen, J., Robinson, J., Ayoglu, B., Mahdessian, D., Sullivan, D., Thul, P., Danielsson, F., Stadler, C., Lundberg, E., Bergstrom, G., Gummesson, A., Voldborg, B. G., Tegel, H., Hober, S., Forsstrom, B., Schwenk, J. M., Fagerberg, L., and Sivertsson, A. (2019) The human secretome. Sci Signal 12

3. Tang, V. T., and Ginsburg, D. (2023) Cargo selection in endoplasmic reticulum-to-Golgi transport and relevant diseases. J Clin Invest 133

4. Zanetti, G., Pahuja, K. B., Studer, S., Shim, S., and Schekman, R. (2012) COPII and the regulation of protein sorting in mammals. Nature Cell Biology 14, 20--28

5. Markova, E. A., and Zanetti, G. (2019) Visualizing membrane trafficking through the electron microscope: Cryo-tomography of coat complexes. Acta Crystallographica Section D: Structural Biology 75, 467--474

6. Weigel, A. V., Chang, C. L., Shtengel, G., Xu, C. S., Hoffman, D. P., Freeman, M., Iyer, N., Aaron, J., Khuon, S., Bogovic, J., Qiu, W., Hess, H. F., and Lippincott-Schwartz, J. (2021) ER-to-Golgi protein delivery through an interwoven, tubular network extending from ER. Cell 184, 2412–2429 e2416

7. Barlowe, C., and Helenius, A. (2016) Cargo Capture and Bulk Flow in the Early Secretory Pathway. Annual Review of Cell and Developmental Biology 32, 197--222

8. Schindler, R., Itin, C., Zerial, M., Lottspeich, F., and Hauri, H. P. (1993) ERGIC-53, a membrane protein of the ER-Golgi intermediate compartment, carries an ER retention motif. Eur J Cell Biol 61, 1–9

9. Nichols, W. C., Seligsohn, U., Zivelin, A., Terry, V. H., Hertel, C. E., Wheatley, M. A., Moussalli, M. J., Hauri, H. P., Ciavarella, N., Kaufman, R. J., and Ginsburg, D. (1998) Mutations in the ER-Golgi intermediate compartment protein ERGIC-53 cause combined deficiency of coagulation factors V and VIII. Cell 93, 61--70

10. Neerman-Arbez, M., Johnson, K. M., Morris, M. A., McVey, J. H., Peyvandi, F., Nichols, W. C., Ginsburg, D., Rossier, C., Antonarakis, S. E., and Tuddenham, E. G. (1999) Molecular analysis of the ERGIC-53 gene in 35 families with combined factor V-factor VIII deficiency. Blood 93, 2253–2260

11. Nyfeler, B., Reiterer, V., Wendeler, M. W., Stefan, E., Zhang, B., Michnick, S. W., and Hauri, H. P. (2008) Identification of ERGIC-53 as an intracellular transport receptor of alpha1-antitrypsin. Journal of Cell Biology 180, 705--712

12. Chen, Y., Hojo, S., Matsumoto, N., and Yamamoto, K. (2013) Regulation of Mac-2BP secretion is mediated by its N-glycan binding to ERGIC-53. Glycobiology 23, 904–916

13. Duellman, T., Burnett, J., Shin, A., and Yang, J. (2015) LMAN1 (ERGIC-53) is a potential carrier protein for matrix metalloproteinase-9 glycoprotein secretion. Biochem Biophys Res Commun 464, 685–691

14. Vollenweider, F., Kappeler, F., Itin, C., and Hauri, H. P. (1998) Mistargeting of the lectin ERGIC-53 to the endoplasmic reticulum of HeLa cells impairs the secretion of a lysosomal enzyme. J Cell Biol 142, 377–389

15. Nyfeler, B., Michnick, S. W., and Hauri, H. P. (2005) Capturing protein interactions in the secretory pathway of living cells. Proc Natl Acad Sci U S A 102, 6350–6355

16. Fu, Y. L., Zhang, B., and Mu, T. W. (2019) LMAN1 (ERGIC-53) promotes trafficking of neuroreceptors. Biochem Biophys Res Commun 511, 356–362

17. Zhang, B., Zheng, C., Zhu, M., Tao, J., Vasievich, M. P., Baines, A., Kim, J., Schekman, R., Kaufman, R. J., and Ginsburg, D. (2011) Mice deficient in LMAN1 exhibit FV and FVIII deficiencies and liver accumulation of alpha1-antitrypsin. Blood 118, 3384–3391

18. Mitrovic, S., Ben-Tekaya, H., Koegler, E., Gruenberg, J., and Hauri, H. P. (2008) The Cargo Receptors Surf4, Endoplasmic Reticulum-Golgi Intermediate Compartment (ERGIC)-53, and p25 Are Required to Maintain the Architecture of ERGIC and Golgi. Molecular Biology of the Cell 19, 1976--1990

19. Armes, N., and Fried, M. (1996) Surfeit locus gene homologs are widely distributed in invertebrate genomes. Mol Cell Biol 16, 5591–5596

20. Belden, W. J. (2001) Role of Erv29p in Collecting Soluble Secretory Proteins into ER-Derived Transport Vesicles. Science 294, 1528--1531

21. Caldwell, S. R., Hill, K. J., and Cooper, A. A. (2001) Degradation of Endoplasmic Reticulum (ER) Quality Control Substrates Requires Transport between the ER and Golgi. Journal of Biological Chemistry 276, 23296--23303

22. Otte, S., and Barlowe, C. (2004) Sorting signals can direct receptor-mediated export of soluble proteins into COPII vesicles. Nature Cell Biology 6, 1189--1194

23. Emmer, B. T., Hesketh, G. G., Kotnik, E., Tang, V. T., Lascuna, P. J., Xiang, J., Gingras, A. C., Chen, X. W., and Ginsburg, D. (2018) The cargo receptor SURF4 promotes the efficient cellular secretion of PCSK9. eLife 7, 1--18

24. Gomez-Navarro, N., Maldutyte, J., Poljak, K., Peak-Chew, S. Y., Orme, J., Bisnett, B. J., Lamb, C. H., Boyce, M., Gianni, D., and Miller, E. A. (2022) Selective inhibition of protein secretion by abrogating receptor-coat interactions during ER export. Proc Natl Acad Sci U S A 119, e2202080119

25. Tang, V. T., McCormick, J., Xu, B., Wang, Y., Fang, H., Wang, X., Siemieniak, D., Khoriaty, R., Emmer, B. T., Chen, X. W., and Ginsburg, D. (2022) Hepatic inactivation of murine Surf4 results in marked reduction in plasma cholesterol. Elife 11

26. Saegusa, K., Sato, M., Morooka, N., Hara, T., and Sato, K. (2018) SFT-4 Surf4 control ER export of soluble cargo proteins and participate in ER exit site organization. Journal of Cell Biology, 1--13

27. Wang, X., Wang, H., Xu, B., Huang, D., Nie, C., Pu, L., Zajac, G. J. M., Yan, H., Zhao, J., Shi, F., Emmer, B. T., Lu, J., Wang, R., Dong, X., Dai, J., Zhou, W., Wang, C., Gao, G., Wang, Y., Willer, C., Lu, X., Zhu, Y., and Chen, X. W. (2021) Receptor-Mediated ER Export of Lipoproteins Controls Lipid Homeostasis in Mice and Humans. Cell Metabolism 33, 350--366.e357

28. Wang, B., Shen, Y., Zhai, L., Xia, X., Gu, H.-m., Wang, M., Zhao, Y., Chang, X., Alabi, A., Xing, S., Deng, S., Liu, B., Wang, G., Qin, S., and Zhang, D.-w. (2021) Atherosclerosis-associated hepatic secretion of VLDL but not PCSK9 is dependent on cargo receptor protein Surf4. Journal of Lipid Research 62, 100091

29. Yin, Y., Garcia, M. R., Novak, A. J., Saunders, A. M., Ank, R. S., Nam, A. S., and Fisher, L. W. (2018) Surf4 (Erv29p) binds amino-terminal tripeptide motifs of soluble cargo proteins with different affinities, enabling prioritization of their exit from the endoplasmic reticulum. PLoS biology 16, e2005140

30. Lin, Z., King, R., Tang, V., Myers, G., Balbin-Cuesta, G., Friedman, A., McGee, B., Desch, K., Ozel, A. B., Siemieniak, D., Reddy, P., Emmer, B., and Khoriaty, R. (2020) The Endoplasmic Reticulum Cargo Receptor SURF4 Facilitates Efficient Erythropoietin Secretion. Molecular and Cellular Biology 40

31. Ordonez, A., Harding, H. P., Marciniak, S. J., and Ron, D. (2021) Cargo receptor-assisted endoplasmic reticulum export of pathogenic alpha1-antitrypsin polymers. Cell Rep 35, 109144

32. Tang, X., Chen, R., Mesias, V. S. D., Wang, T., Wang, Y., Poljak, K., Fan, X., Miao, H., Hu, J., Zhang, L., Huang, J., Yao, S., Miller, E. A., and Guo, Y. (2022) A SURF4-to-proteoglycan relay mechanism that mediates the sorting and secretion of a tagged variant of sonic hedgehog. Proc Natl Acad Sci U S A 119, e2113991119

33. Saegusa, K., Matsunaga, K., Maeda, M., Saito, K., Izumi, T., and Sato, K. (2022) Cargo receptor Surf4 regulates endoplasmic reticulum export of proinsulin in pancreatic beta-cells. Commun Biol 5, 458

34. Devireddy, S., and Ferguson, S. M. (2022) Efficient progranulin exit from the ER requires its interaction with prosaposin, a Surf4 cargo. J Cell Biol 221

35. Tang, X., Wang, T., and Guo, Y. (2022) Export of polybasic motif-containing secretory proteins BMP8A and SFRP1 from the endoplasmic reticulum is regulated by surfeit locus protein 4 (SURF4): SURF4 regulates ER export of BMP8A and SFRP1. J Biol Chem, 102687

36. Huang, Y., Yin, H., Li, B., Wu, Q., Liu, Y., Poljak, K., Maldutyte, J., Tang, X., Wang, M., Wu, Z., Miller, E. A., Jiang, L., Yao, Z.-P., and Guo, Y. (2021) An in vitro vesicle formation assay reveals cargo clients and factors that mediate vesicular trafficking. Proceedings of the National Academy of Sciences 118, e2101287118

37. Grube, L., Dellen, R., Kruse, F., Schwender, H., Sthler, K., and Poschmann, G. (2018) Mining the Secretome of C2C12 Muscle Cells: Data Dependent Experimental Approach to Analyze Protein Secretion Using Label-Free Quantification and Peptide Based Analysis. Journal of Proteome Research 17, 879--890

38. Poschmann, G., Brenig, K., Lenz, T., and Stuhler, K. (2021) Comparative Secretomics Gives Access to High Confident Secretome Data: Evaluation of Different Methods for the Determination of Bona Fide Secreted Proteins. Proteomics 21, e2000178

39. Abbineni, P. S., Tang, V. T., da Veiga Leprevost, F., Basrur, V., Xiang, J., Nesvizhskii, A. I., and Ginsburg, D. (2022) Identification of secreted proteins by comparison of protein abundance in conditioned media and cell lysates. Anal Biochem 655, 114846

40. Nakabayashi, H., Miyano, K., Sato, J., Yamane, T., and Taketa, K. (1982) Growth of human hepatoma cell lines with differentiated functions in chemically defined medium. Cancer Research 42, 3858--3863

41. Sanjana, N. E., Shalem, O., and Zhang, F. (2014) Improved vectors and genome-wide libraries for CRISPR screening. Nat Methods 11, 783–784

42. Emmer, B. T., Hesketh, G. G., Kotnik, E., Tang, V. T., Lascuna, P. J., Xiang, J., Gingras, A. C., Chen, X. W., and Ginsburg, D. (2018) The cargo receptor SURF4 promotes the efficient cellular secretion of PCSK9. Elife 7

43. Purushothaman, A. (2019) Exosomes from Cell Culture-Conditioned Medium: Isolation by Ultracentrifugation and Characterization. Methods Mol Biol 1952, 233–244

44. da Veiga Leprevost, F., Haynes, S. E., Avtonomov, D. M., Chang, H. Y., Shanmugam, A. K., Mellacheruvu, D., Kong, A. T., and Nesvizhskii, A. I. (2020) Philosopher: a versatile toolkit for shotgun proteomics data analysis. Nat Methods 17, 869–870

45. Kong, A. T., Leprevost, F. V., Avtonomov, D. M., Mellacheruvu, D., and Nesvizhskii, A. I. (2017) MSFragger: ultrafast and comprehensive peptide identification in mass spectrometry-based proteomics. Nat Methods 14, 513–520

46. Djomehri, S. I., Gonzalez, M. E., da Veiga Leprevost, F., Tekula, S. R., Chang, H. Y., White, M. J., Cimino-Mathews, A., Burman, B., Basrur, V., Argani, P., Nesvizhskii, A. I., and Kleer, C. G. (2020) Quantitative proteomic landscape of metaplastic breast carcinoma pathological subtypes and their relationship to triple-negative tumors. Nat Commun 11, 1723

47. UniProt, C. (2021) UniProt: the universal protein knowledgebase in 2021. Nucleic Acids Res 49, D480–D489

48. Ritchie, M. E., Phipson, B., Wu, D., Hu, Y., Law, C. W., Shi, W., and Smyth, G. K. (2015) limma powers differential expression analyses for RNA-sequencing and microarray studies. Nucleic Acids Res 43, e47

49. Nyfeler, B., Zhang, B., Ginsburg, D., Kaufman, R. J., and Hauri, H. P. (2006) Cargo selectivity of the ERGIC-53/MCFD2 transport receptor complex. Traffic 7, 1473–1481

50. Zhang, B., Zheng, C., Zhu, M., Tao, J., Vasievich, M. P., Baines, A., Kim, J., Schekman, R., Kaufman, R. J., and Ginsburg, D. (2011) Mice deficient in LMAN1 exhibit FV and FVIII deficiencies and liver accumulation of alpha1-antitrypsin. Blood 118, 3384--3391

51. Shen, Y., Wang, B., Deng, S., Zhai, L., mei Gu, H., Alabi, A., Xia, X., Zhao, Y., Chang, X., Qin, S., and wei Zhang, D. (2020) Surf4 regulates expression of proprotein convertase subtilisin/kexin type 9 (PCSK9) but is not required for PCSK9 secretion in cultured human hepatocytes. Biochimica et Biophysica Acta (BBA) - Molecular and Cell Biology of Lipids 1865, 158555

52. Zhang, B., McGee, B., Yamaoka, J. S., Guglielmone, H., Downes, K. A., Minoldo, S., Jarchum, G., Peyvandi, F., de Bosch, N. B., Ruiz-Saez, A., Chatelain, B., Olpinski, M., Bockenstedt, P., Sperl, W., Kaufman, R. J., Nichols, W. C., Tuddenham, E. G., and Ginsburg, D. (2006) Combined deficiency of factor V and factor VIII is due to mutations in either LMAN1 or MCFD2. Blood 107, 1903–1907

53. Emmer, B. T., Lascuna, P. J., Kotnik, E. N., Saunders, T. L., Khoriaty, R., and Ginsburg, D. (2019) Murine Surf4 is essential for early embryonic development. bioRxiv

54. Yan, R., Chen, K., Wang, B., and Xu, K. (2022) SURF4-induced tubular ERGIC selectively expedites ER-to-Golgi transport. Dev Cell 57, 512–525 e518

